# UROPA GUI: A web platform for genomic region annotation

**DOI:** 10.1101/302091

**Authors:** Hendrik Schultheis, Jens Preussner, Annika Fust, Mette Bentsen, Carsten Kuenne, Mario Looso

## Abstract

The annotation of genomic ranges such as peaks resulting from ChIP-seq/ATAC-seq or other techniques represents a fundamental task of bioinformatics analysis with considerable impact on many downstream analyses. In our previous work, we introduced the Universal Robust Peak Annotator (UROPA), a flexible command line based tool which improves upon the functionality of existing annotation software. In order to reduce the complexity for biologists and clinicians, we have implemented an intuitive web-based graphical user interface (GUI) and fully functional service platform for UROPA. This extension will empower all users to generate annotations for regions of interest interactively.

**Availability and Implementation:** The open source UROPA GUI server was implemented in R Shiny and Python and is available from http://loosolab.mpi-bn.mpg.de. The source code of our App can be downloaded at https://github.molgen.mpg.de/loosolab/UROPA_GUI under the MIT license.

## Introduction

Annotation in the context of peaks deriving from e.g. ChIP-seq/ATAC-seq data denotes the identification of relevant genomic features such as transcripts, genes, or promoters in range. Reference feature loci can derive from public databases (e.g. NCBI RefSeq, Ensembl/Gencode) or custom reference data sets. The attribution of meaningful features to a peak is mandatory to permit the mapping to existing databases bearing transcript or gene identifiers which are needed for downstream analyses such as pathway enrichment, ontology assignments, and others.

Popular tools addressing peak annotation tasks include Homer [1], GREAT [2], Goldmine [3], and CHIPpeakAnno [4], covering a number of use cases ranging from simple annotations with limited options (e.g. Homer feature assignment is based on the closest NCBI RefSeq transcription start site (TSS)) to complex annotations with numerous parameters (e.g. Goldmine can integrate all features sets available from the UCSC Table Browser). However all of them rely on command line tools, and require a certain level of computational skills. GREAT additionally provides a web-based graphical user interface (GUI), but is limited by the number of dated feature reference sets and only supports a fixed set of association rules. A user friendly GUI, able to provide an advanced annotation algorithm with a large variety of parameters to an audience of biologists and clinicians with limited computational skills is critically missing.

Our previously introduced command line based tool **U**niversal **Ro**bust **P**eak **A**nnotator (UROPA) comprises a wide range of unique features to individually adapt the annotation process to the experimental setup [5]. UROPA is not only intended to provide feature assignments based on classical gene start/end models, but also offers functionality for relative positioning of BED formatted genomic regions of interest with respect to any GTF formatted feature set given by the user. While arguably simpler to use than competing command line tools, proper configuration of the association rules for UROPA requires some experience. Furthermore, UROPA relies on a local Linux operating system including Python and R configured with the necessary packages.

Here, we introduce a user friendly, intuitive and web-based graphical user interface for UROPA, providing all functionality of UROPA to non-computational scientists. UROPA GUI is the first web application for peak annotation, which can integrate multiple user defined association rules (=queries) including optional prioritization. These rules can be defined in a step by step process based on a predefined vocabulary, thus eliminating a range of prevalent causes for error when compared to manual configuration on the command line. The public web server includes a variety of feature reference data sets available from Ensembl and permits the upload of custom reference GTF files.

## Using the UROPA GUI Server

The web-based UROPA GUI was implemented using R Shiny, internally calling an updated UROPA Python script. The workflow and interface of the UROPA GUI server is presented in Figure 1A. Starting from genomic regions of interest provided by the user as a BED formatted file, the desired reference feature set can either be selected from existing data on the server or can be uploaded via a file upload dialog. The server provides pre-calculated gene annotation sets for 14 widely used model organisms. Based on the features parsed from the selected reference file, the user is asked to define at least one association rule (query) that defines which features will be selected for annotation. An optional global priority flag defines if the order of the generated queries has to be taken into account. The GUI provides an interactive form, granting granular access to all optional parameters that can be addressed by UROPA with respect to each single location of interest, such as distance to feature, orientation (upstream/downstream), biotype, and others explained in more detail in the manual (http://uropa-manual.readthedocs.io/). By submitting the query via the **Run_UROPA** button, the job is queued to a cluster and the browser session is redirected to a result page that will automatically update after successful job completion, and that allows subsequent download of results. Multiple output files are generated to cover various granularities of assigned features per genomic region: 1) all assigned candidate features from each query, 2) the best assigned feature from each query, and 3) the best assigned feature among all queries. Furthermore, a set of graphs is generated in order to visualize the annotation efficiency and the contribution of single queries to overall assignment rates, provided as a PDF. Each web session receives a unique UROPA identifier (ID), intended for temporary persistence of results. Since all computational load of UROPA is handled by a performant cluster, there is no need for users to install UROPA locally or to provide computing power on premise. Notably, the UROPA GUI web server is open source and can be used without user registration or login. The source code for UROPA and UROPA_GUI is freely available from our GitHub repository for offline usage respectively (https://github.molgen.mpg.de/loosolab/UROPA, https://github.molgen.mpg.de/loosolab/UROPA_GUI).

**Figure 1.**
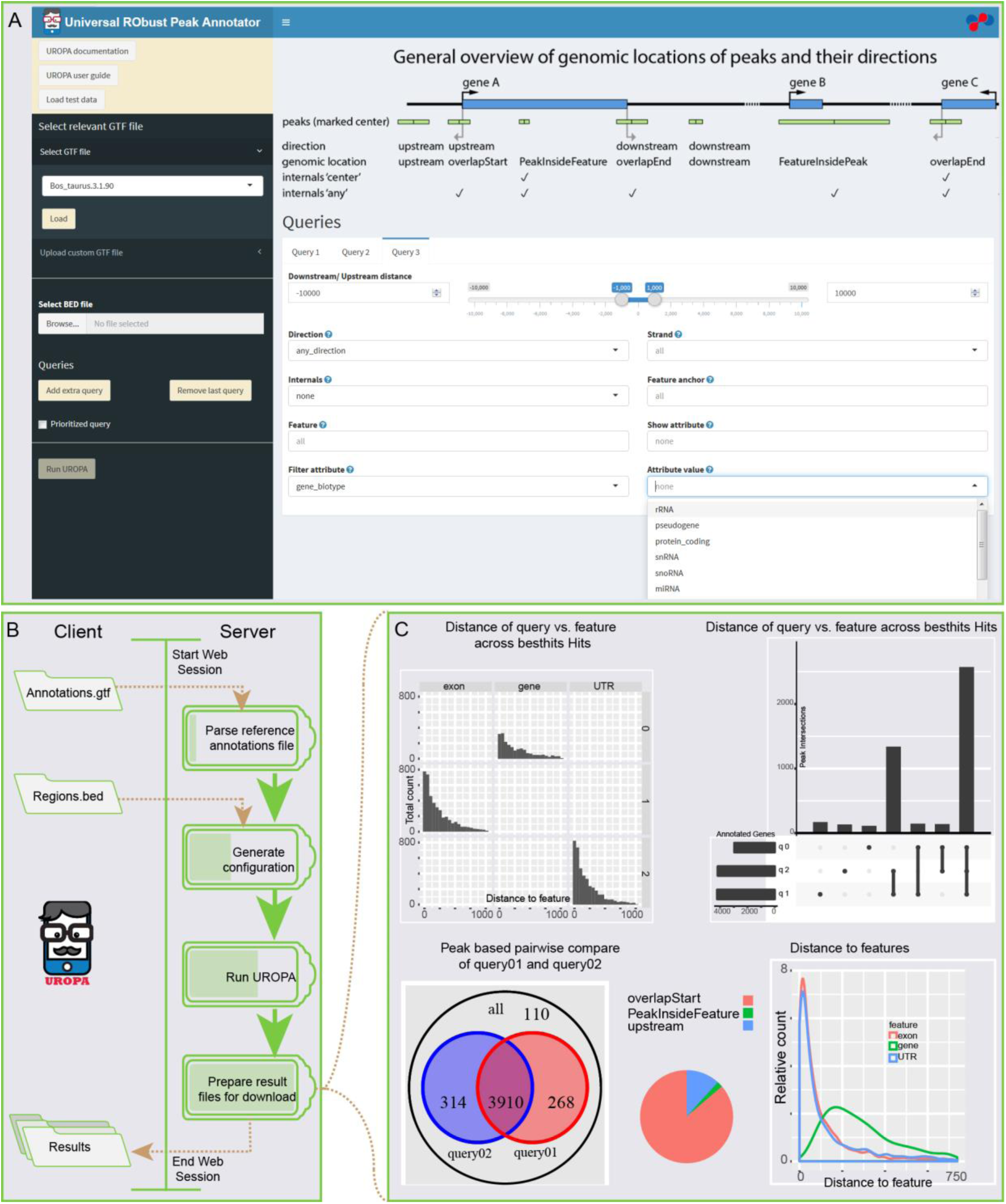
Workflow and appearance of UROPA GUI; A) appearance of the UROPA GUI with upload and file selection dialogs on the left (black) and a detailed query design field on the right (Query board). B) dataflow (brown dashed lines) and workflow (green) between a web client that provides genomic regions of interest and the UROPA GUI server. C) Exemplary output visualizations of an successful UROPA run, as provided for download from the UROPA GUI server.

## Authors’ contributions

HS, JP, CK and ML conceived the algorithm, HS, JP, MB and AF implemented the algorithm, CK set up web services and infrastructure. CK, JP and ML wrote the manuscript. All authors read and approved it.

## Acknowledgements and Funding

This work was supported by the Max-Planck Society, the Excellence Cluster Cardio-Pulmonary System (ECCPS), and from the Deutsche Forschungsgemeinschaft (KLIFO309). We thank Peter Hofmann, Franz Ziegengeist and Hosro Lotfikhah for IT support.

## References

1. Heinz S, Benner C, Spann N, Bertolino E, Lin YC, Laslo P, et al. Simple combinations of lineage-determining transcription factors prime cis-regulatory elements required for macrophage and B cell identities. Mol Cell. 2010;38(4):576-89.

2. McLean CY, Bristor D, Hiller M, Clarke SL, Schaar BT, Lowe CB, et al. GREAT improves functional interpretation of cis-regulatory regions. Nat Biotechnol. 2010;28(5):495-501.

3. Bhasin JM, Ting AH. Goldmine integrates information placing genomic ranges into meaningful biological contexts. Nucleic Acids Res. 2016;44(12):5550-6.

4. Zhu LJ, Gazin C, Lawson ND, Pages H, Lin SM, Lapointe DS, et al. ChIPpeakAnno: a Bioconductor package to annotate ChIP-seq and ChIP-chip data. BMC Bioinformatics. 2010;11:237.

5. Kondili M, Fust A, Preussner J, Kuenne C, Braun T, Looso M. UROPA: a tool for Universal RObust Peak Annotation. Sci Rep. 2017;7(1):2593.

